# Microbial Diversity and Ecological Structure of the Gut Microbiota in Children and Adults from an Indigenous Mexican Community

**DOI:** 10.1101/2025.08.03.668344

**Authors:** Sebastián Villada-Bedoya, Luisa I. Falcón, Osiris Gaona, Aída Elizondo García, Isaac G-Santoyo

**Affiliations:** Neuroecology Lab, Facultad de Psicología, UNAM, Mexico City 04510, Mexico; Laboratorio de Ecología Bacteriana, Instituto de Ecología, UNAM, Mexico City 04510, Mexico; Instituto de Ecología, Campus Yucatán, Parque Científico y Tecnológico de Yucatán, Mérida 97302, Mexico

**Keywords:** gut bacteria microbiota, human health, human microbial diversity, indigenous lifestyle

## Abstract

Gut microbiota composition is shaped by environmental factors such as lifestyle, diet, socioeconomic and cultural context, as well as host-related factors like age and sex. Although microbiota research has expanded rapidly in recent years, most current paradigms are based on data from individuals in high-income, industrialized countries, particularly in Europe and North America. As a result, microbiota profiles from non-Westernized populations—especially those from Indigenous communities—remain underrepresented in global studies, limiting our understanding of the full ecological and evolutionary diversity of human gut microbial communities. In this study, we characterized the bacterial gut microbiota of Me’phaa individuals, a pre-Columbian Indigenous group from the south-central region of México. This rural community maintains a subsistence lifestyle centered on high-fiber, plant-based diets and limited exposure to industrialized foods. Using high-throughput 16S rRNA gene sequencing, we analyzed microbial profiles from 62 individuals aged 5 to 60 years, examining differences across age and sex to explore patterns of diversity and community structure. Despite sharing similar life conditions, noticeable differences in gut microbiota were observed; alpha and beta diversities revealed significant variation across age and sex. Children exhibited greater microbial richness and inter-individual variability, consistent with dynamic colonization processes that begin in early life. In contrast, adults showed more taxonomically uniform communities, with a higher abundance of taxa such as Prevotellaceae, Succinivibrio, and Dialister, which are associated with fiber fermentation and complex carbohydrate metabolism. Additionally, sex-related differences were evident, with adult males displaying the highest relative abundance of Prevotellaceae. Finally, the presence of taxonomic groups rarely observed in urban Western populations suggests that indigenous lifestyles may preserve microbial diversity with potential metabolic relevance. Our findings advance the ecological and evolutionary understanding of human gut microbiota and highlight the importance of integrating non-Westernized populations into microbiome research to better elucidate host-microbe coadaptation and its implications for health.

## Introduction

The human body harbors a vast number of microorganisms, most of which reside in the gut and collectively are referred to as the gut microbiota (GM) (1). Among these microorganisms, bacteria represents the most diverse group, with ∼3180 genera described to date (2). The GM plays a crucial role in regulating key host biological host processes (3) that impact its health (4–6), including immunity, nutrient metabolism, and neuronal functioning (7–10). The host-microbiota relationship is essentially symbiotic: the host provides nutrients and ecological niches, while microorganisms contribute metabolic, protective, and immunomodulatory functions that the host cannot perform alone (11,12). Hence, alterations in this host-symbiont relationships have been observed in several human pathologies such as obesity, type II diabetes mellitus, bowel irritable bowel syndrome, cardiometabolic disorders, among others (13).

Lifestyle practices play a critical role in shaping the structural ecology of the gut microbiota (14,15). Key factors include mode of delivery (cesarean vs. vaginal birth), infant feeding practices, access to allopathic medicine, and long-term dietary habits (10). Another key determinant is the host’s age. It has been reported that the rapid bacterial diversification observed in early childhood slows between the first and fifth years of life (16). Between the ages of seven and twelve, the gut microbiota becomes progressively more similar to that of adults (17–19). However, some evidence suggests that microbial communities may still retain taxonomic and functional distinctions during this stage (20–22).

Despite the rapid growth and dynamism of microbiome research, most current paradigms regarding lifestyle-associated variation are based on data from populations residing in high-income countries in Europe and North America, grouped under what is commonly referred to as “Westernized lifestyles.” Limited efforts have been made to explore gut microbial ecology in populations whose lifestyles fall outside this category, including hunter-gatherers such as the Hadza of Tanzania (23,24), subsistence farmers in Bassa, Nigeria (25), indigenous agriculturalists in Bangladesh (3), farmers in Papua New Guinea (26), the agropastoral Wiwa community in Colombia (27), Amerindian groups in South America (28,29), and the Inuit from the Canadian Arctic (10).

From these studies, it is now understood that such populations harbor a GM with greater phylogenetic diversity compared to Westernized groups (26,28–30). Additionally, these populations—regardless of continent—are enriched in a recurrent set of bacterial groups collectively known as VANISH taxa (Volatile or Associated Negatively with Industrialized Societies of Humans), including the families Prevotellaceae, Spirochaetaceae, and Succinivibrionaceae. These taxa are specialized in degrading complex plant-derived, fiber-rich carbohydrate due to their high abundance of carbohydrate-active enzymes (CAZymes) (14,24). In contrast, they show minimal or no presence of BIoSSUM taxa (Bloom or Selected in Urbanization/Modernization Societies), which include members of the Bacteroidaceae, Enterobacteriaceae, and Verrucomicrobiaceae families (14,30).

Despite this progress, a large gap remains in our understanding of the gut microbial ecology of other world regions. For instance, aside from the studies in Venezuelan Amazonian Amerindian groups (29), very little is known about non-Westernized populations in Latin America’s Global South—particularly those representing pre-Hispanic indigenous communities. Currently, 826 indigenous groups are officially recognized across Latin America (31), encompassing between 42 and 50 million individuals (32) dispersed over approximately 20.1 million km² of territory. Due to a long history of colonialism, indigenous groups from Latin America’s Global South exhibits a wide spectrum of lifestyle practices, ranging from Westernized behaviors to highly pre-Columbian modes of living reflecting a profound ethnocultural mosaic, with populations at the farthest extremes of what is practiced in industrialized societies.

In Mexico alone, at least 78 indigenous groups are recognized (31), whose lifestyles span this entire spectrum—from highly Westernized to minimally Westernized. One example is the Me’phaa people of the Montaña Alta region in Guerrero, an indigenous group residing in isolated communities of 50–80 families (each composed of five to ten members) (33,34). Their subsistence economy is based on the cultivation of maize and legumes (e.g., beans and lentils), collection of wild edible plants, and occasional fruit and vegetable farming. Animal protein is sourced through hunting and poultry farming, usually reserved for special occasions (33). All food is locally grown and harvested nearby (35). These communities lack sanitation infrastructure and potable water, relying on small wells, and have minimal access to allopathic medicine; vaginal births account for 98% of deliveries, and breastfeeding continues until at least two years of age. Residents speak only their native language, face extremely low income levels, and experience high infant and adult morbidity and mortality rates due to their geographic and social isolation (36).

Thus, the aim of this study was to characterize the population structure, diversity, and taxonomic composition of the bacterial gut microbiota of a Me’phaa indigenous community in south-central Mexico, and to identify potential associations with age and sex. To this end, we addressed the following research questions: i) How do taxonomic assignments and diversity indices differ between age and sex groups within the Me’phaa community? and ii) Can age or sex predict higher or lower abundances of VANISH and BIoSSUM taxa in this non-Westernized population? We hypothesized that total microbial diversity would be lower in children compared to adults.

Additionally, we anticipated a high prevalence of VANISH taxa—linked to fiber-rich diets—and low or absent representation of BIoSSUM taxa typically found in Westernized microbiomes.

The study of this population offers an opportunity to understand how sociocultural practices shape gut microbial ecology in non-Westernized Latin American populations. Moreover, they allow us to compare findings not only with the few existing studies of other non-Westernized groups around the world but also with Westernized populations whose lifestyles may overlap in specific dimensions (e.g., fiber-rich diets yet high exposure to modern medicine). Ultimately, these insights expand our understanding of global health by offering a less biased view of human microbiome variation.

## Materials and methods

### Data collection

The present study was conducted in two Me’phaa communities, Plan de Gatica (17° 7’44.364“ N, 99°7’22.695” W; 510 m a.s.l.) and El Naranjo (17°9’54.0036” N, 98°57’50.9832” W; 860 m a.s.l.) in the state of Guerrero, Mexico. The collection of biological material were conducted from September 7 to 29, 2017 and consisted of human fecal samples from children and adults belonging to Both indigenous communities are located approximately 30 km apart and share similar socioeconomic and cultural patterns (35). The diet in these communities is low in fat and animal protein and is characterized by the consumption of plant-derived carbohydrates from a fiber- and cereal-rich diet (14,37). The inclusion and exclusion criteria for this study were as follows: i) children aged 5–12 and adults aged 18–46, ii) born in the Plan de Gatica and Naranjo community and having lived there all their lives, iii) not having taken any type of allopathic medicine in the last 6 months or any type of antibiotic in the last 2 years, iv) in children, all born by natural childbirth and having been breastfed during early childhood and at least until the first 2 years of life, and v) not reporting any type of gastrointestinal or respiratory condition two months prior to sample collection.

Fecal samples were obtained from children (aged between 5 and 10 years) and adults (aged between 19 and 76 years) in the Me’phaa community. A total of 62 fecal samples were collected, comprising 33 adults (15 males and 18 females) and 29 children (13 males and 16 females). Although the small sample size may be a limitation, other authors have worked with similar sample sizes and obtained robust results (25,38). Each participant was provided with a kit containing a sterile plastic container and a wooden tongue depressor to facilitate sample collection. Each container was labeled with a unique code assigned to each participant. Once the samples were returned by participants, they were processed: three aliquots of approximately 100 µL of fecal material were prepared using sterile pipette tips and transferred into sterile 1.5 mL microtubes. All fecal samples were frozen in liquid nitrogen and stored at -20 °C until DNA extraction.

In addition to fecal samples, the study included anthropometric measurements (i.e., height, weight, BMI) for all participants, as well as a questionnaire on the nutritional status of both children and adults. Furthermore, information was collected regarding pregnancy, delivery, and breastfeeding duration. For the Me’phaa community, assistance from a translator was required, as this community does not speak Spanish.

### Fecal DNA extraction

DNA was extracted from each of the 62 fecal samples using the DNeasy Blood & Tissue Kit (Qiagen, Valencia, CA, USA), following the manufacturer’s instructions. Each sample (∼100 µL of fecal DNA) was resuspended in 30 µL of molecular-grade water and stored at -20 °C until PCR amplification.

### 16S rRNA gene amplification and sequencing

Using the previously extracted DNA, the hypervariable V4 region of the 16S rRNA gene was amplified with universal bacterial/archaeal primers 515F/806R (14,39), following the procedures described by Caporaso et al. (2012) and Sánchez-Quinto et al. (2020) (38,40). PCR reactions were performed in triplicate for each sample, in a final volume of 25 µL, prepared as follows (reagents mixed in this order): 1) 15.67 µL of molecular-grade nuclease-free water, 2) 2.5 µL of Takara ExTaq 10× PCR buffer, 3) 0.7 µL of bovine serum albumin (BSA, 20 mg mL⁻¹), 4) 2 µL of Takara dNTP mix (2.5 mM), 5) 1 µL of 515F primer (10 µM), 6) 0.125 µL of Takara ExTaq DNA polymerase (5 U µL⁻¹) (TaKaRa, Shiga, Japan), 7) 2 µL of DNA template, and 8) 1 µL of 806R primer.

The PCR protocol included an initial denaturation step at 95 °C for 3 minutes, followed by 35 cycles of 95 °C for 30 seconds, 52 °C for 40 seconds, and 72 °C for 90 seconds, with a final extension at 72 °C for 12 minutes. Once amplification was complete for all samples, triplicate reactions were pooled and purified using SPRI magnetic beads with the Agencourt AMPure XP PCR purification system (Beckman Coulter, Brea, CA, USA). The purified DNA was quantified using a Qubit 2.0 fluorometer to obtain approximately 20 ng per sample.

The purified 16S rRNA amplicons (∼20 ng) were sequenced on an Illumina MiSeq platform at the Yale Center for Genome Analysis (New Haven, CT, USA), generating paired-end reads of approximately 250 bp. All sequences obtained were uploaded to the NCBI database under BioProject accession number PRJNA593240.

### Analysis of the sequence data

Paired-end 2 × 250 reads were processed in QIIME2 and denoised using the DADA2 plugin to resolve amplicon sequence variants (ASVs) (41). Forward and reverse reads were truncated at 200 bp, and chimeric sequences were removed using the “consensus” method. Taxonomic assignment of representative ASV sequences was performed using the “classify consensus-vsearch” plugin (42), using the SILVA 132 database as a reference (43). Sequence alignment was performed using the MAFFT algorithm (44). After masking conserved positions and filtering gaps, a phylogeny was constructed with the FastTree algorithm (45). The abundance table and the phylogeny were exported into the R environment to perform the statistical analyses using phyloseq (46) and ggplot2 (47) packages. Plastic ASVs were filtered from the samples, which were then rarefied to a minimum sequencing of 21,000 reads per sample. A Venn diagram showing ASV distribution by age group was generated using the ‘ps_venn’ function from the MicEco package in R (48). Total diversity (α diversity) of ASVs was calculated using Faith’s phylogenetic diversity (PD) index, Shannon diversity index, Simpson’s dominance index, and the observed ASVs.

### Statistical analysis

All statistical analyses were performed in R version 4.3.0 (49) using RStudio (50). For alpha (α) diversity analysis, observed richness (observed ASVs), Shannon index, inverse Simpson index, and Faith’s PD were calculated to assess overall richness and diversity of the sequenced samples across age groups and sexes. Statistical differences between these factors were determined using Type III analysis of variance tables (GLM; generalized linear models) with the ‘Anova’ function from the car package (51). Data were summarized and organized using the dplyr package (52), and figures were generated with ggplot2 (47). Beta (β) diversity analysis between age groups and sexes was estimated by calculating weighted UniFrac and Bray-Curtis distances. Statistical differences were assessed through permutational multivariate analysis of variance (PERMANOVA) using distance matrices. Additionally, a linear discriminant analysis (LDA) effect size (LEfSe) was performed. This algorithm identifies and explains high-dimensional biomarkers, detecting genomic features (genes, pathways, or taxa) that characterize differences between two or more biological conditions or classes. This analysis emphasizes statistical significance, biological consistency, and effect relevance, allowing identification of differentially abundant features that are also consistent with biologically meaningful categories (53).

Moreover, differences in the abundance of VANISH families (Prevotellaceae, Spirochaetaceae, and Succinivibrionaceae) and BIoSSUM families (Bacteroidaceae, Enterobacteriaceae, and Verrucomicrobiaceae) (14) between age groups and sexes were explored using GLMs or zero-inflated regression models (ZAGA), as appropriate, implemented with the gamlss package (54).

### Ethics

All procedures for testing and recruitment were approved (25 September 2017) by the National Autonomous University of Mexico Committee on Research Ethics (FPSI/CE/01/2016) and run in accordance with the ethical principles and guidelines of the Mexican Law (NOM-012-SSA3-2012). All participants read and signed a written informed consent. Additionally, we received signed approval from all participants (or their parents) who were photographed for the manuscript.

## Results

### Taxonomic comparison of the GM between indigenous Me’phaa children and adults

This study generated a dataset of 62 fecal samples, corresponding to 29 children aged 5 to 10 years (16 girls and 13 boys) and 33 adults aged 19 to 76 years (18 females and 15 males) from the Me’phaa indigenous group (high mountain region of Guerrero). A total of 1,302,000 sequences were recovered after quality filtering and chimera removal. Firmicutes and Bacteroidota were the dominant phylum in both children and adults (Fig 1a). Within each age group, Firmicutes was the dominant phylum, showing the highest relative abundance (females = 68.86%, males = 52.15%, girls = 76.73%, and boys = 74.78%), followed by Bacteroidota (females = 22.10%, males = 40.46%, girls = 13.97%, and boys = 13.00%). A general pattern was observed in which children (both girls and boys) exhibited substantially lower relative abundances of Bacteroidota compared to adults.

**Fig 1.**
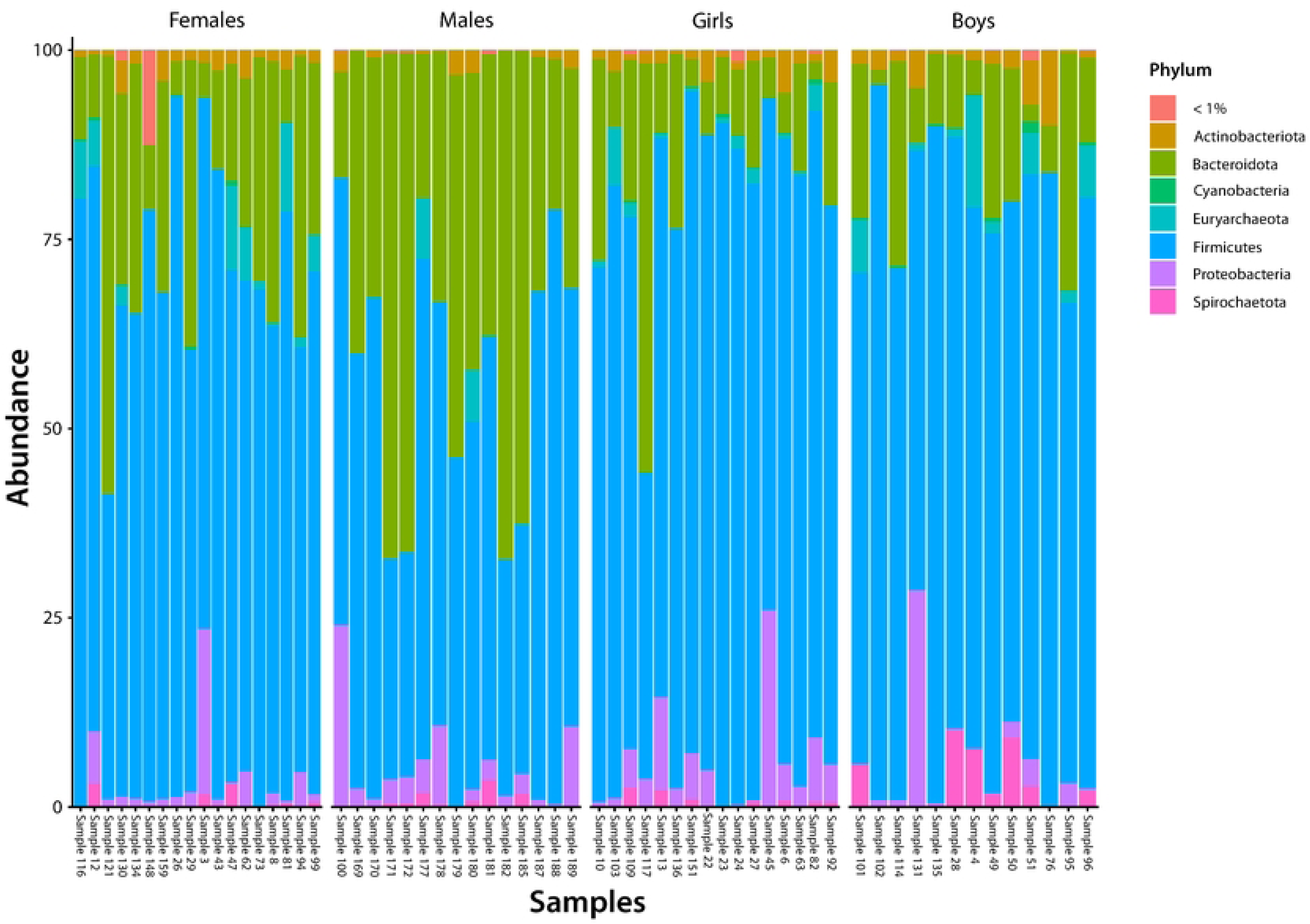
**(a)** Relative abundance of the gut microbiota and distribution of bacterial composition (16S rRNA V4) at the Phylum level in children and adults from the Me’phaa community. Phyla with relative abundances < 1% were grouped into the “< 1%” category. **(b)** Venn diagram of ASVs from the Me’phaa community.

The phylum Proteobacteria showed considerable relative abundances in males (4.61%) and girls (5.22%). Conversely, the phylum Euryarchaetota displayed higher relative abundances in females (2.98%) and boys (3.01%), while Spirochaetota was more abundant in boys (2.99%) (Fig 1a). Some phyla were exclusively present in children (both boys and girls), including Chloroflexi (0.028%), Deinococcota (0.017%), Myxococcota (0.014%), Planctomycetota (0.013%), and Gemmatimonadota (0.008%). Moreover, the Venn diagram (Fig 1b) revealed that half (50.05%) of the total ASVs were shared between age groups. The ASVs specific to adults and children (25.28% and 24.67%, respectively) indicated that shared diversity was greater than the diversity specific to each age group (Fig 1b). This same pattern persisted when further discriminated by both age group and sex, where the shared diversity was substantially higher (ASVs = 50.86%) than the ASVs specific to each subgroup (females = 11.56%, males = 11.01%, girls = 10.31%, and boys = 10.26%).

At the family level, the dominant families were Ruminococcaceae, Prevotellaceae, and Lachnospiraceae in both children and adults (S1 Fig). Within each age group, Ruminococcaceae was the dominant family, showing the highest relative abundance (females = 20.91%, males = 17.64%, girls = 21.25%, and boys = 22.05%). Notably, in females, Prevotellaceae exhibited the highest relative abundance (38.26%), whereas in the other age groups this family was the second most abundant (females = 18.57%, girls = 11.69%, and boys = 11.73%). Lachnospiraceae was the third most abundant family (females = 13.99%, males = 12.05%, girls = 12.54%, and boys = 13.60%). It was observed that the adult population (both females and males) had higher relative abundances of Prevotellaceae compared to children.

The family Erysipelatoclostridiaceae showed considerable relative abundances in females (4.59%) and boys (5.66%). Conversely, the families Mycoplasmataceae, Streptococcaceae, and Enterobacteriaceae exhibited notable relative abundances in girls (9.15%, 5.20%, and 4.04%, respectively) (S1 Fig). Some families were exclusive to children (both boys and girls), including Woeseiaceae (0.173%), Xenococcaceae (0.476%), Hydrogenophilaceae (0.409%), Nitrosococcaceae (0.034%), and Desulfosarcinaceae (0.305%).

### Alpha and beta diversity of the GM in the indigenous Me’phaa populations

When evaluating the observed ASVs in children and adults from the Me’phaa community, statistically significant differences were found for the interaction between age group × sex (Table 1; Fig 2a). Boys exhibited greater richness in their GM, with these differences primarily observed between boys and girls (t = 2.655, df = 54, *P* = 0.04) and between boys and females (t = 2.902, df = 54, *P* = 0.02).

**Fig 2.**
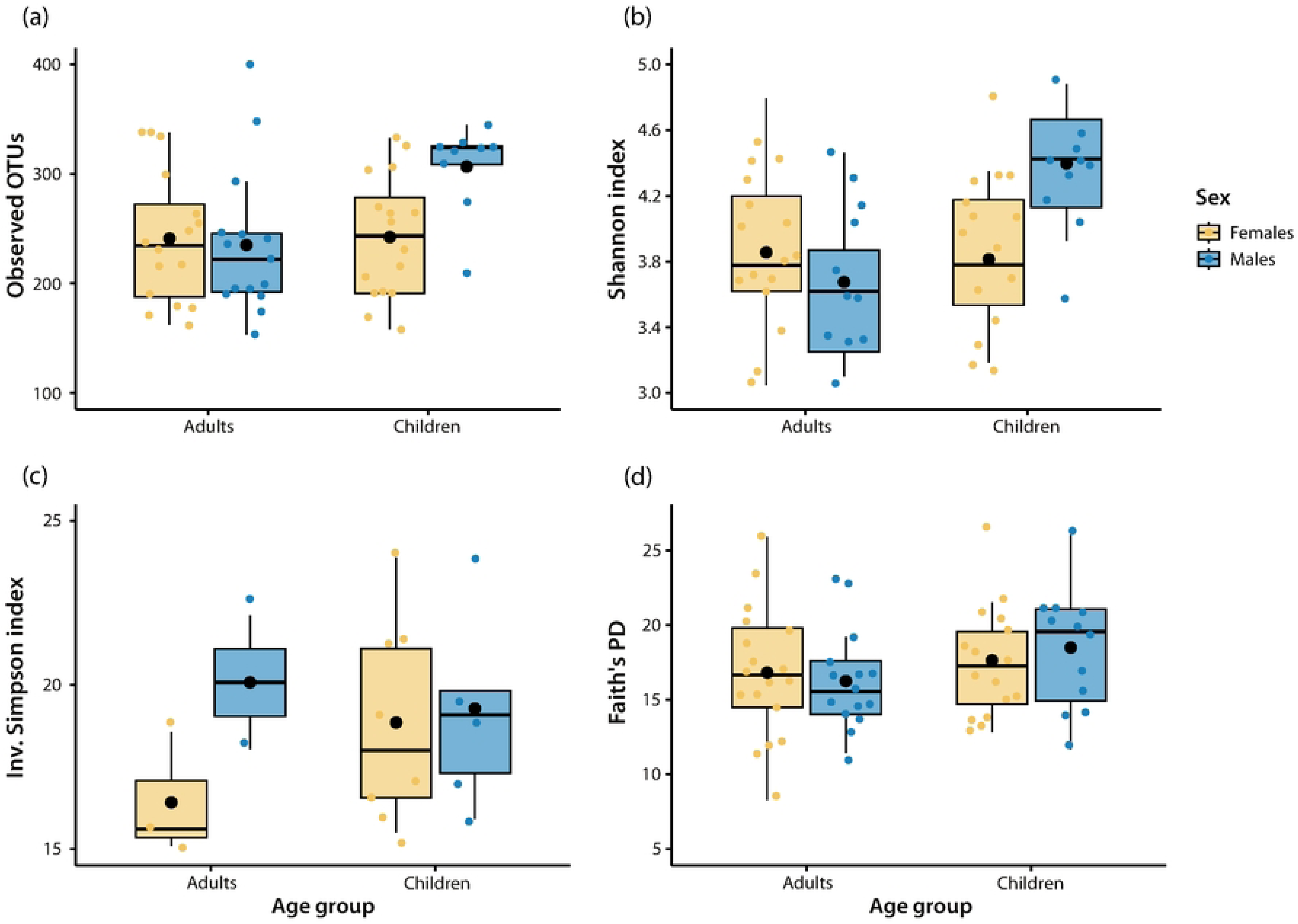
Alpha diversity indices of the gut microbiota in Me’phaa populations by age group and sex. **(a)** Observed OTUs; **(b)** Shannon index; **(c)** Simpson dominance index (inverse); **(d)** Faith’s phylogenetic diversity index. The line inside each box represents the median; boxes represent the first and third quartiles, and whiskers indicate the maximum and minimum values (excluding outliers). The black dot represents the mean. Each colored dot represents an individual/participant. Significant differences (*P* < 0.05; see Table 1).

In terms of the Shannon diversity index, statistically significant differences were found for age group and for the age group × sex interaction (Table 1; Fig 2b). Again, boys showed higher diversity values in their gut microbiota, with significant differences mainly between boys and girls (t = -3.358, df = 53, *P* < 0.01), between boys and males (t = -4.048, df = 53, *P* < 0.001), and between boys and females (t = -3.446, df = 53, *P* < 0.01).

The Simpson dominance index indicated statistically significant differences for both explanatory variables and their interaction (Table 1). These differences in dominant ASVs were mainly observed between boys and males (t = -2.864, df = 57, *P* = 0.02) and between boys and females (t = -2.774, df = 57, *P* = 0.03) (Fig 2c). Finally, for Faith’s PD index, only marginally significant differences were found between age groups, specifically between children and adults of both sexes (t = 1.801, df = 58, *P* = 0.08) (Table 1; Fig 2d).

**Table 1.**
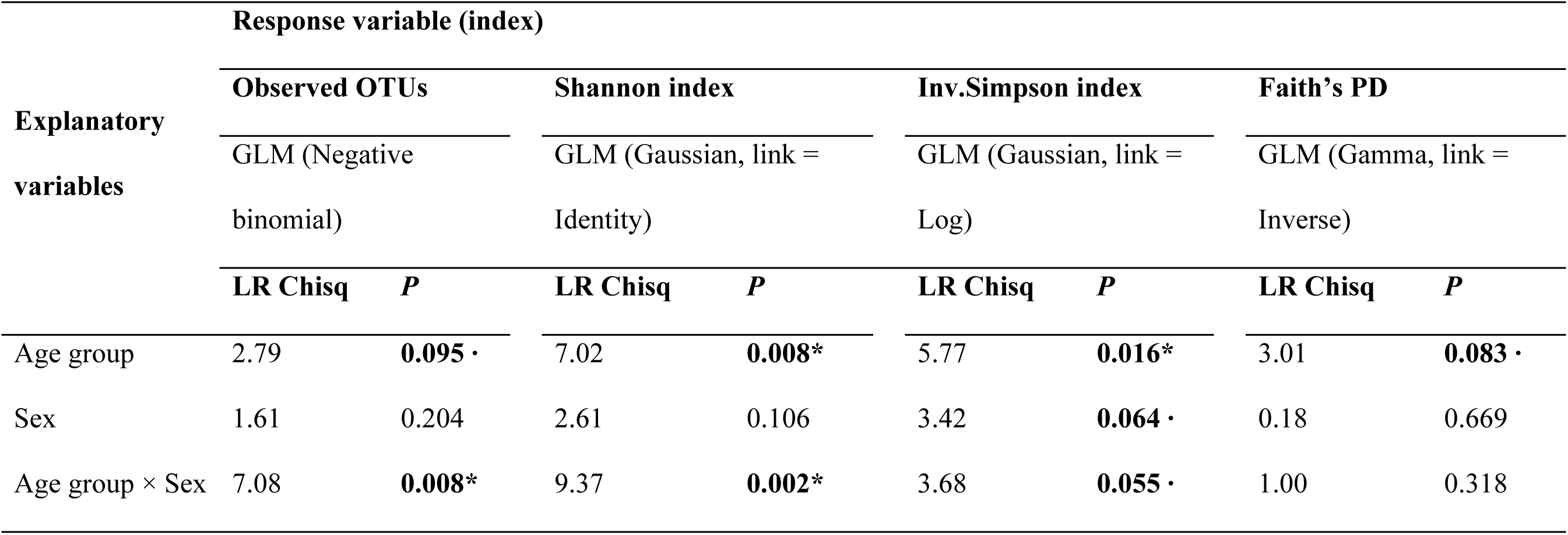
Results of the generalized linear models (GLM) for each diversity index evaluated. LR Chisq = likelihood ratio Chi-square. (**\*)** Significant differences; (**·**) Marginally significant differences, both indicated in bold.

When evaluating beta diversity of the gut microbiota in children and adults from the Me’phaa community, no clear separation between age groups was evident. Using the weighted UniFrac method, marginally significant differences were found for the age group × sex interaction (PERMANOVA, F = 2.25, R² = 0.04, *P* = 0.080); children exhibited greater dispersion, while adults showed tighter clustering (Fig 3a). Using the Bray–Curtis method, the age group × sex interaction showed statistically significant differences (PERMANOVA, F = 2.26, R² = 0.04, *P* = 0.001), and a separation between children and adults was observed (Fig 3b), suggesting that the composition of the gut microbiota may vary by age group and sex in these indigenous populations.

**Fig 3.**
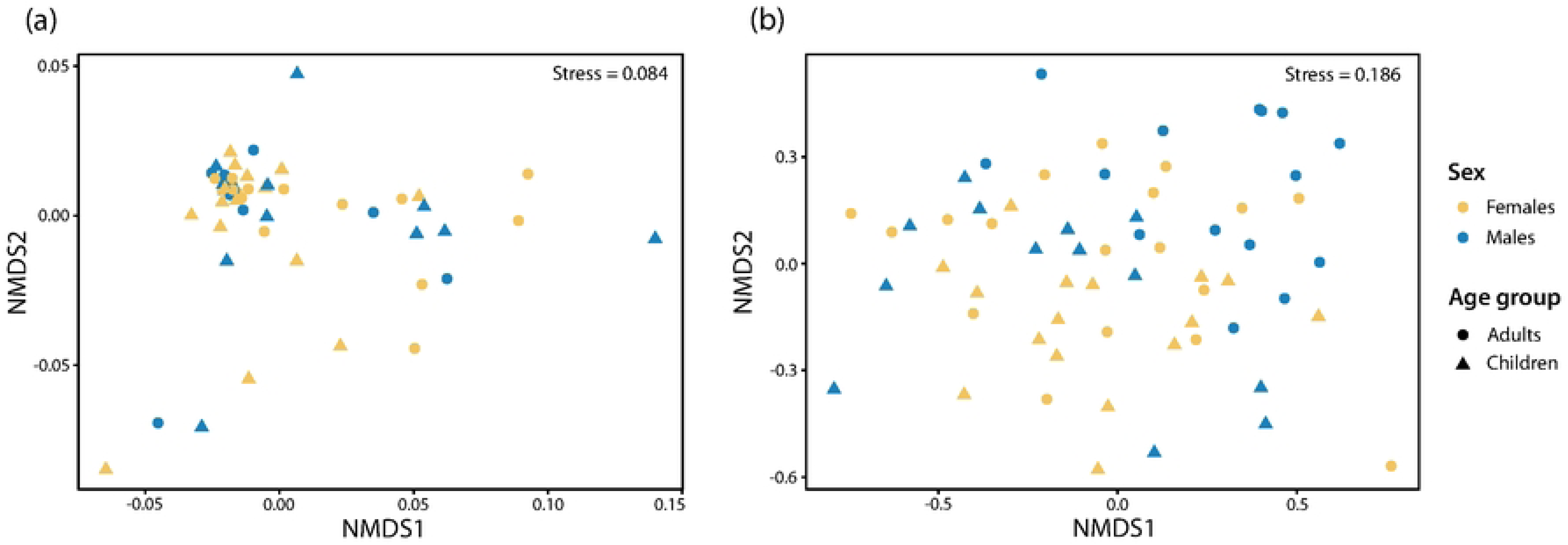
Non-metric multidimensional scaling (NMDS) analysis showing β diversity of Me’phaa indigenous populations. **(a)** Clustering based on the Weighted UniFrac method. **(b)** Clustering based on the Bray–Curtis index. Statistically significant differences were detected using PERMANOVA at a significance level of *P* < 0.05.

Both distance metrics were independent of sex (PERMANOVA, weighted UniFrac: F = 0.64, R² = 0.01, *P* = 0.61; Bray–Curtis: F = 0.91, R² = 0.01, *P* = 0.52). Intergroup differences in gut microbiota composition between children and adults from the Me’phaa community were evident and are reflected in taxonomic differences between these two age groups. The heatmap of the most abundant ASVs illustrates similarities and differences between Me’phaa children and adults, as well as between sexes (Fig 4), with a higher prevalence of the Prevotellaceae family in adults and genera such as *Actinomyces* and *Akkermansia* in children.

**Fig 4.**
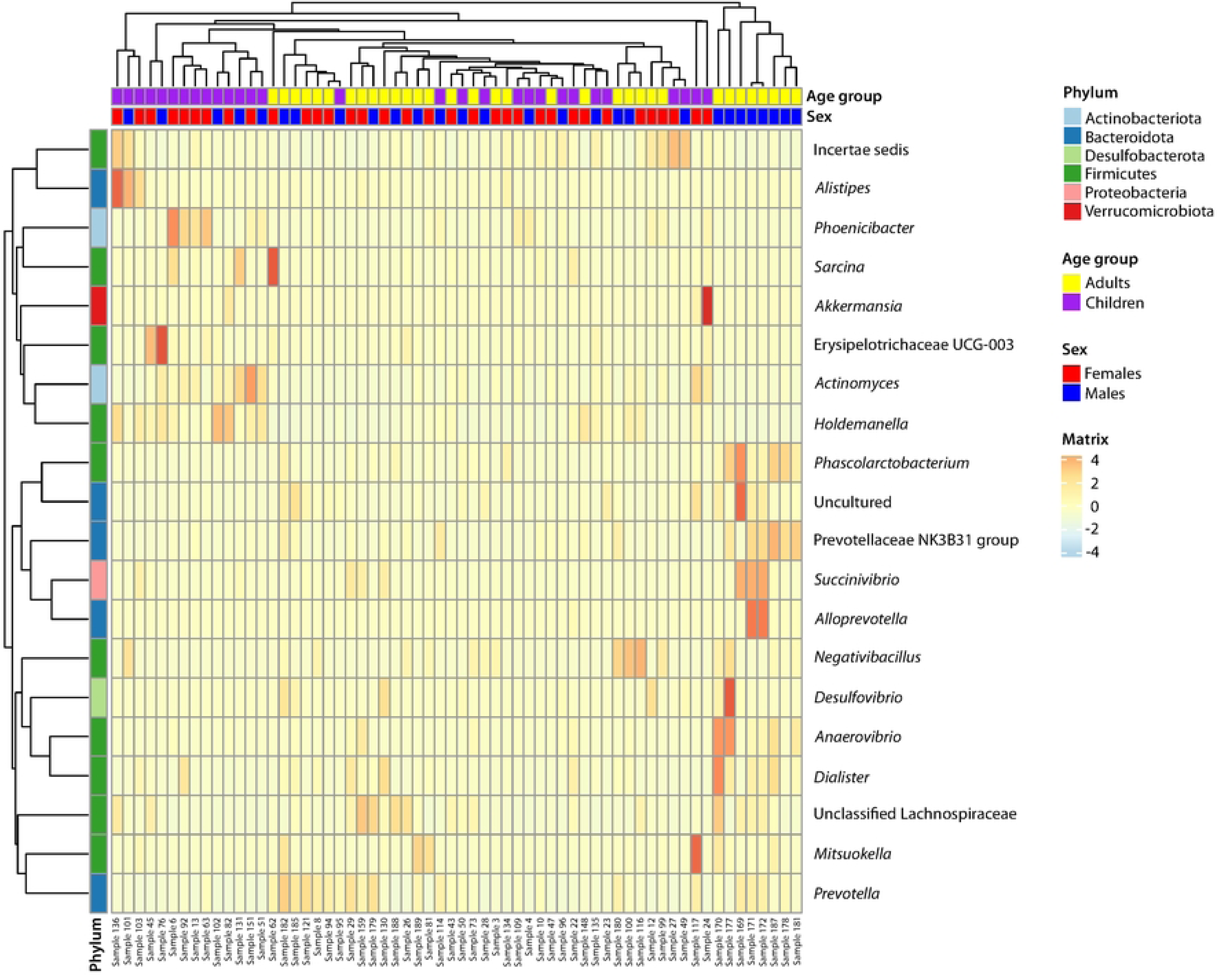
Heatmap of the most abundant ASVs in Me’phaa children and adults. The left side of the plot displays the ASV clustering dendrogram, and the top shows the sample clustering dendrogram. Each color in the central heatmap grid represents the relative abundance value of ASVs in each taxon after normalization processing.

### Analysis of differences in the GM between Me’phaa indigenous children and adults

To further investigate taxonomic distributions and clades in Me’phaa children and adults, we compared microbial community abundances at each taxonomic level. The histogram from the linear discriminant analysis (LDA) effect size (LEfSe) shows the LDA scores for microbial taxa with significant differences across age groups and sexes, indicating that females exhibited fewer microbial taxa compared to other groups (Fig 5). Conversely, the LEfSe cladogram visualizes all detected microbial communities (abundance >1%) from phylum to genus (Fig 6). Based on the inferred taxonomic profiles across all samples, LEfSe performed a statistical strategy to discover metagenomic biomarkers and identified 50 differentially abundant taxa (S1 Table).

**Fig 5.**
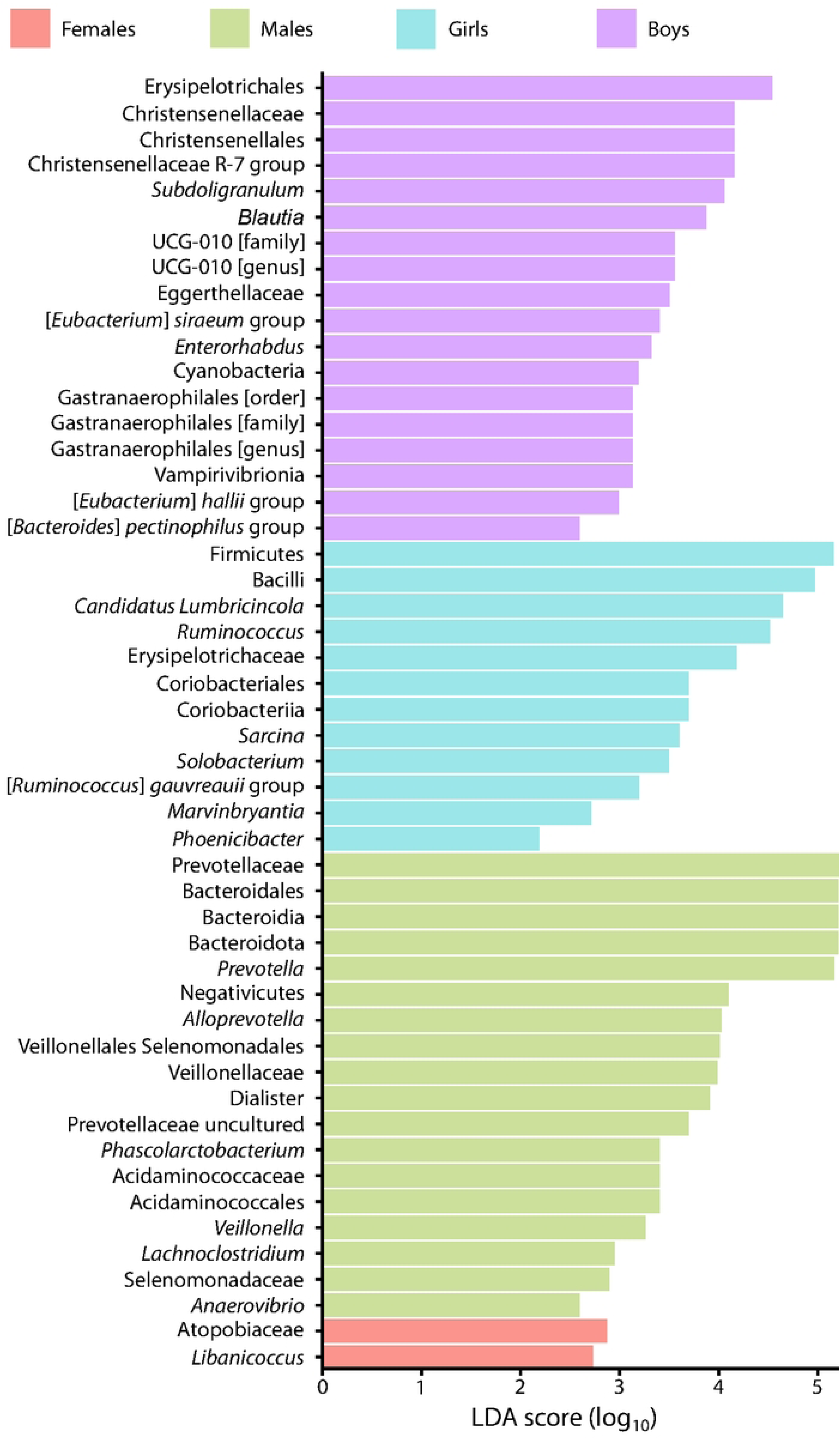
Histogram of the LEfSe analysis. The length of each bar represents the LDA score. The figure shows microbial taxa with significant differences across age groups and sexes. Females (red), males (green), girls (blue), and boys (purple). LDA score > 2.

**Fig 6.**
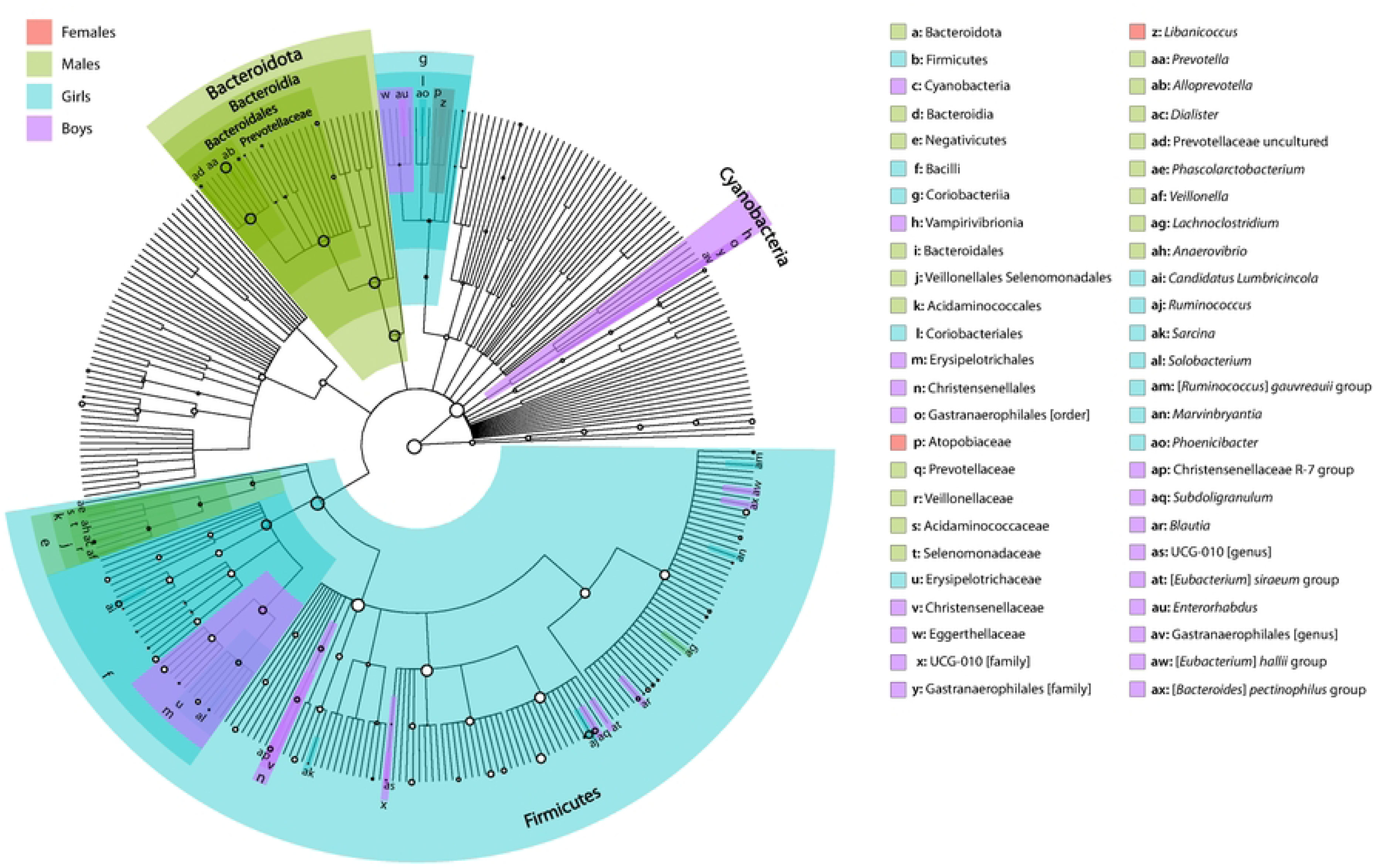
Cladogram from the LEfSe analysis, showing differences in the relative abundance of taxa at five levels (Phylum, Class, Order, Family, and Genus) between Me’phaa children and adults. Circles of different colors indicate that age groups and sexes exhibited differences in relative abundance. White circles represent non-significant taxa.

Taxa that best characterized the females group included the family Atopobiaceae and the genus *Libanicoccus*. For males, the most representative taxa were from the families Prevotellaceae and Veillonellaceae, and the genera *Alloprevotella* and *Dialister*. In the children’s age group, the most abundant microbial taxa in girls belonged to the families Mycoplasmataceae and Ruminococcaceae, and the genera *Candidatus Lumbricincola* (a new lineage) and *Ruminococcus*, respectively. Finally, in boys, the most dominant families were Christensenellaceae, Ruminococcaceae, and Lachnospiraceae, with the group Christensenellaceae (R-7) and the genera *Subdoligranulum* and *Blautia*, respectively (S1 Table; Fig 6).

### Differences between VANISH and BIoSSUM groups in Me’phaa indigenous children and adults

There were differences in the abundance of taxa belonging to VANISH groups (volatile and/or negatively associated with industrialized human societies) and BIoSSUM groups (positively associated with urban/modernized societies) (14). For this reason, in this study, we considered age group and sex of Me’phaa indigenous children and adults as predictors for the following families: Prevotellaceae, Spirochaetaceae, and Succinivibrionaceae (VANISH), and Bacteroidaceae, Enterobacteriaceae, and Verrucomicrobiaceae (BIoSSUM). It is worth noting that although the population studied is an indigenous community, and therefore BIoSSUM-associated taxa are likely not highly represented, we decided to carry out analyses for both groups.

The GM of Me’phaa indigenous children and adults showed significant differences in two out of the three VANISH groups (Table 2). For Prevotellaceae, significant differences were found for all explanatory variables, as well as for the age group × sex interaction (Table 2), where males exhibited higher abundance compared to the other groups (Fig 7a). For the Spirochaetaceae family, there were significant differences for the age group × sex interaction (Table 2), with boys showing the highest abundance compared to the other groups (Fig 7b). In the case of the Succinivibrionaceae family, no significant differences were detected for any of the variables or interactions evaluated (Table 2; Fig 7c).

**Table 2.**
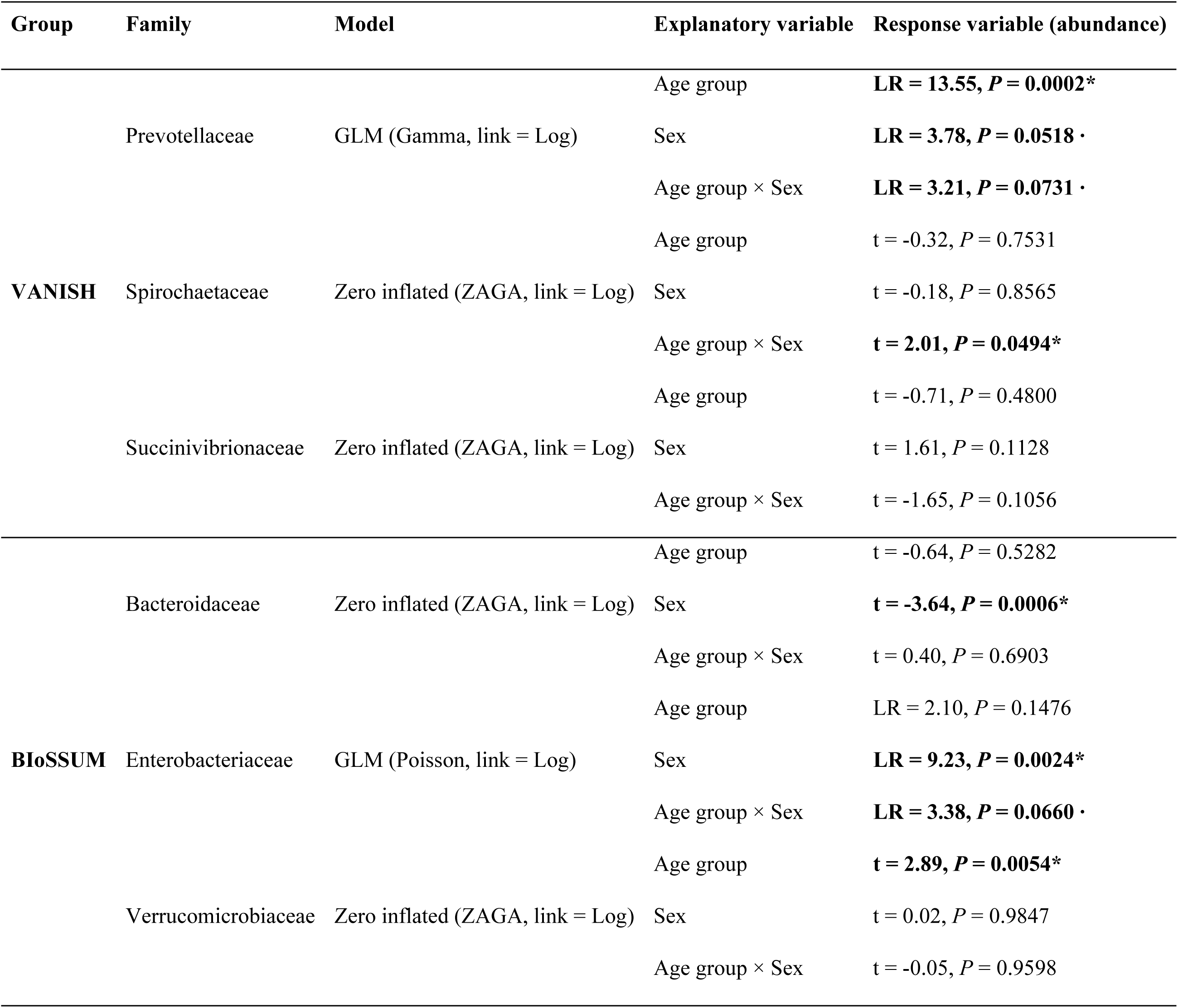
Results of the linear models for each family belonging to the VANISH and BIoSSUM groups. LR Chisq = likelihood ratio chi-squared statistic. t = t-test. **(*)** Significant differences; **(·)** Marginally significant differences, both shown in bold.

**Fig 7.**
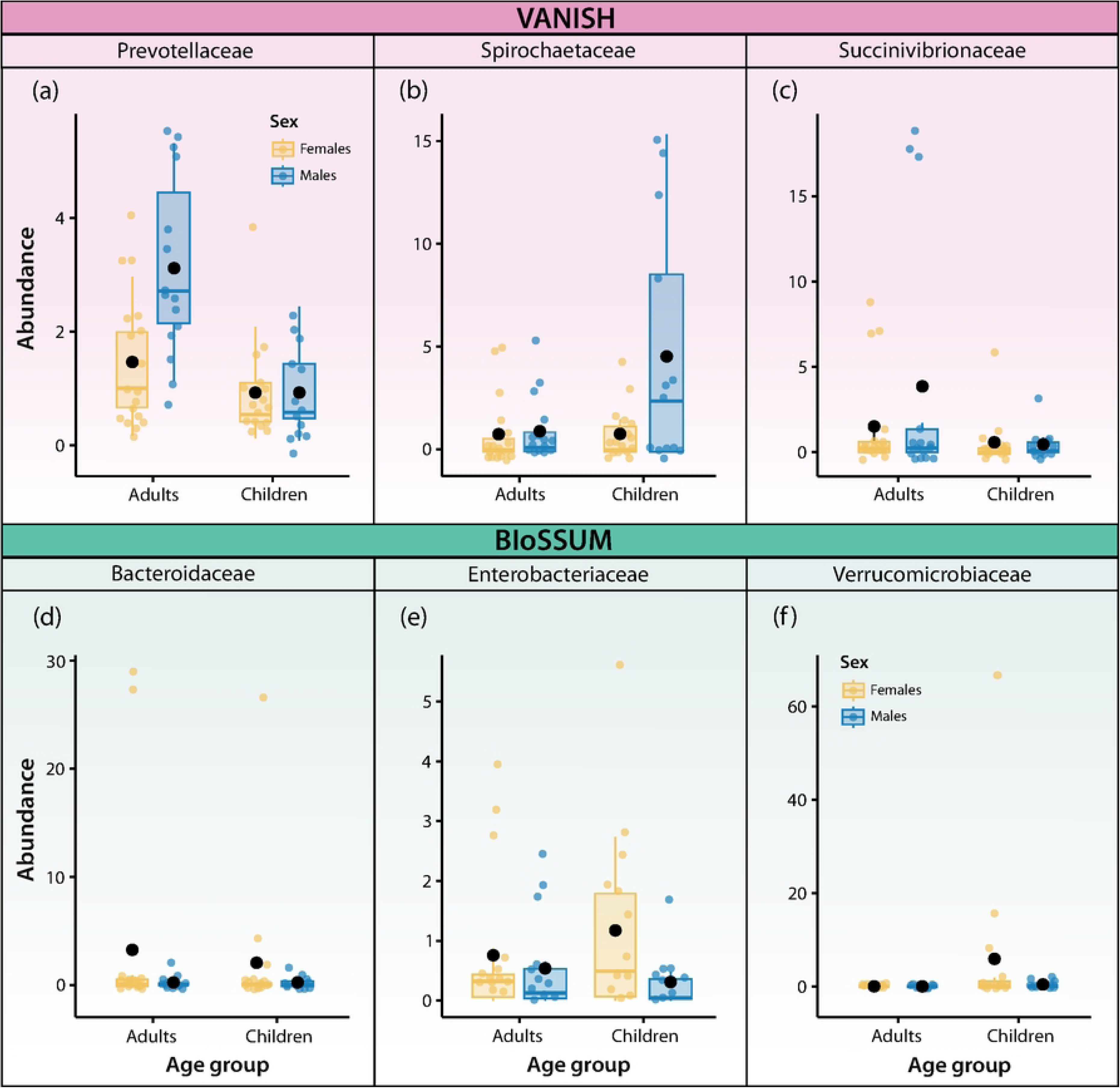
Boxplots of VANISH and BIoSSUM GM groups in Me’phaa indigenous children and adults. The plots show family-level taxa in the fecal microbiota of Me’phaa children and adults. The line inside each box represents the median, the boxes represent the first and third quartiles, and the whiskers indicate minimum and maximum values. The black dot represents the mean. Each colored dot represents an individual/participant. Significant differences (*P* < 0.05; see Table 2).

Regarding families in the BIoSSUM group, significant differences were observed across all three families (Table 2). For Bacteroidaceae, significant differences were found between sexes (Table 2), with females exhibiting higher abundances than males, regardless of age group (Fig 7d). For the Enterobacteriaceae family, significant differences were observed between sexes, and marginally significant differences for the age group × sex interaction (Table 2), with girls showing the highest abundances (Fig 7e). Finally, for the Verrucomicrobiaceae family, significant differences were observed between age groups, regardless of sex (Table 2), with children exhibiting higher abundances than adults (Fig 7f).

## Discussion

This study contributes to the limited body of knowledge regarding the GM of non-Westernized indigenous populations in the global south of Latin America. We observed that the Mexican Me’phaa population exhibits a remarkably diverse bacterial microbiota, which is consistent with findings in other non-Westernized populations across different geographic regions (10,27). This high diversity is mainly associated with a diet rich in plant-derived carbohydrates, particularly dietary fiber and cereals (37,38). Thus, our findings support the hypothesis that geography or host genetics may play a limited role in defining the microbial ecology of the human gut (55). Furthermore, the study provides insights into how the lifestyle and diet of the Me’phaa communities, contribute to the high diversity and unique microbial composition of their inhabitants.

The second key finding here was the variation observed across participant age groups. In Westernized American populations, gut microbial diversity in children under 3 years is typically lower than in adults (16), and reaches a climax around 3–5 years of age, stabilizing thereafter, including into adulthood (56). In contrast, in this study, alpha diversity tended to be higher in boys compared to girls and adults (both male and female), even though the age range for the children (5– 12 years) is often underrepresented in microbiota studies, given the presumed stability reached by that age. Interestingly, the most common phyla among non-Westernized populations include Firmicutes, Bacteroidetes, Actinobacteria, and Verrucomicrobia (57). In Me’phaa children, we also observed representation from Actinobacteriota, Cyanobacteria, Euryarchaeota, Proteobacteria, and Spirochaetota (Fig. 1a). Notably, some phyla were detected exclusively in children (both boys and girls), albeit at very low abundances (0.008–0.028%), including Chloroflexi, Deinococcota, Myxococcota, Planctomycetota, and Gemmatimonadota. These groups, although uncommon in the human GM, have been reported in other non-Westernized populations and may reflect increased exposure to environmental microorganisms (e.g., soil, untreated water, and plant-based materials), as well as diets rich in plant fibers, characteristic of rural communities (29,38). The phylum Chloroflexi represents a low-abundance component (<1%) and remains poorly understood in terms of its physiology, host adaptation, or pathogenic potential (58). For Planctomycetota, some studies in Senegalese populations have reported antimicrobial activity, anaerobic fermentation capacity, and the ability to degrade various polysaccharides and glycoproteins (59). Deinococcota, Myxococcota, and Gemmatimonadota are rare taxa with little or no known human presence, yet their detection may be significant, as low-abundance taxa are thought to play crucial roles in maintaining gut homeostasis and function (57).

We also observed in Me’phaa children a strong presence of the family Christensenellaceae, known for its high heritability and association with gut ecology and diseases such as obesity and IBD (60). The genera *Actinomyces* and *Holdemanella* were dominant among children of both sexes. Girls showed higher levels of Mycoplasmataceae and Ruminococcaceae, and the genera *Ruminococcus*, *Akkermansia*, *Alistipes*, and *Phoenicibacter*. In boys, Christensenellaceae, Ruminococcaceae, and Lachnospiraceae were more abundant, along with the genera *Subdoligranulum* and *Blautia*. *Akkermansia* has been associated with reduced weight gain and fat accumulation, similar to Christensenellaceae (60), and has the capacity to degrade intestinal mucin (61). It also improves glucose tolerance and reduces inflammation and metabolic endotoxemia in animal models (61). Although both taxa share similar metabolic and inflammatory roles, their long-term health effects on the host remain to be fully understood (38).

The lower alpha diversity among adults may reflect a partial transition toward a more Westernized diet (10), characterized by reduced consumption of fiber-rich foods and increased intake of Western dietary components (14). This shift may be explained by the fact that adult men are often the ones who temporarily leave their communities to seek work in more urbanized or densely populated areas. Although short-term dietary changes can influence GM composition, evidence suggests that in healthy individuals, the bacterial profile tends to remain stable and is more significantly shaped by long-term dietary patterns (62–64). In adults from this community, the family Prevotellaceae and the genera *Prevotella*, *Succinivibrio*, and *Dialister* were predominant, consistent with fiber-rich diets and the fermentation of complex polysaccharides (65,66). *Prevotella* is a key biomarker of plant-based diets and has been linked to both metabolic protection and intestinal inflammation (67). Its abundance is associated with high intake of non-digestible polysaccharides due to its robust fermentative capabilities (68), whereas its loss correlates with reduced dietary fiber consumption (14). Short-term consumption of an animal-protein-rich diet has been shown to significantly alter the gut microbiota, increasing the abundance of *Bacteroides*, *Bilophila*, and *Alistipes*, while reducing fiber-degrading Firmicutes (15).

Among the diverse gut microbiota, certain genera such as *Coprococcus* and *Faecalibacterium* have been associated with improved quality of life, largely through the production of butyrate, a short-chain fatty acid (SCFA) that helps reduce intestinal inflammation (69,70). Both genera tend to be depleted in individuals with inflammatory bowel disease (IBD) (71). *Coprococcus* has also been linked to hypertension and autism spectrum disorders (72,73), with sex-specific differences in prevalence (74). *Dialister* appears more abundant in males than females, and in adults compared to children, potentially reflecting greater dietary exposure to Western foods, especially among adult males. *Dialister* is a producer of SCFAs such as propionate and has been associated with immunomodulatory pathways and gut health (75,76). Overall, the elevated presence of these taxa suggests that adults with a pre-Columbian Mexican lifestyle harbor gut microbial profiles specialized in plant compound fermentation, potentially offering metabolic and immunological benefits.

The GM is influenced by lifestyle, diet, ethnicity (15,77,78), and medical practices such as excessive antibiotic use (79). While diet is a major driver of GM structure, non-Westernized populations are often more vulnerable to malnutrition. When it occurs during breastfeeding or early childhood, it can lead to reduced levels of key gut bacteria such as *Bifidobacterium longum*, an important species associated with human milk (13,80). Here, we found low representation of this taxon in both children and adults.

On the other hand, because VANISH taxa are rare in Western populations (on which most studies have been conducted), little is known about their role in the gut (14). Interestingly, the families Spirochaetaceae and Succinivibrionaceae contain highly motile members; however, the functions of these motile taxa within the microbiota remain largely unknown (14). The high abundance of *Prevotella*, a VANISH group, among Me’phaa males (and to a lesser extent in boys), is consistent with findings by Sánchez-Quinto et al. (2020) and other studies conducted in non-Westernized groups such as the Yanomami (Amerindians) and the BaAka (Africans) (29,81).

In the human gastrointestinal tract, symbiotic relationships occur between the GM and the host, through which a large number of peptides are synthesized, surpassing those encoded by the human genome (89). The relevance of this interaction lies in its involvement in essential functions such as vitamin biosynthesis, polysaccharide fermentation, ion absorption, and the regulation of host metabolic pathways (82,83). It also contributes to the production of: i) metabolites such as folate (vitamin B9), indoles, secondary bile acids, and trimethylamine-N-oxide (linked to cardiovascular risk); ii) neurotransmitters such as serotonin and GABA; and viii) SCFAs, critical for gut health (84). These compounds play roles in immune, digestive, metabolic, and central nervous system (CNS) processes (76,84). The GM also contributes to fiber and starch catabolism, pathogen defense (85–88), and gut-brain axis communication, influencing motivation, stress response, and mood disorders (76,89,90).

In conclusion, individuals from the Me’phaa community display high gut microbial diversity, exclusive low-abundance taxa, and clear age- and sex-related differences. The strong representation of bacterial families uncommon in Western populations reinforces the view that non-Westernized lifestyles preserve complex microbiotas with unexplored ecological and health-relevant functions. Future studies should include metagenomic and metabolomic analyses to characterize these functions and assess how lifestyle transitions influence gut microbial structure and function. Indigenous populations offer a unique opportunity to document and conserve human microbial diversity in the context of accelerating globalization and dietary change in the global south of the Americas.

## Acknowledgments

We are grateful to Margarita Muciño, Julio Naranjo, and Diego Hernández-Muciño from Xuajin Me’Phaa, a non-governmental association, for their help in the liaison with the Me’phaa community, and their help with sample collection and logistics. We also are grateful to Olga A. Rojas-Ramos, Ariatna Hernández Castillo, and Rosa M. de la Fuente Rodríguez for their contributions in sample collection. To Daniel Cerqueda-García for his help in bioinformatic analysis. We also thank to UNAM Posdoctoral Program (POSDOC), because during S.V.-B.’s postdoctoral stay it was possible to carry out this work in parallel to his stay in the Neuroecology Lab.

## Author contributions

**Conceptualization:** Isaac G.-Santoyo, Luisa I. Falcón.

**Data curation:** Sebastián Villada-Bedoya, Luisa I. Falcón, Osiris Gaona, Aída Elizondo García, Isaac G.-Santoyo.

**Formal analysis:** Sebastián Villada-Bedoya, Osiris Gaona, Luisa I. Falcón.

**Funding acquisition:** Isaac G.-Santoyo.

**Investigation:** Isaac G.-Santoyo, Luisa I. Falcón, Sebastián Villada-Bedoya, Osiris Gaona, Aída Elizondo García.

**Methodology:** Sebastián Villada-Bedoya, Luisa I. Falcón, Osiris Gaona, Aída Elizondo García, Isaac G.-Santoyo.

**Project administration:** Isaac G.-Santoyo.

**Resources:** Isaac G.-Santoyo, Luisa I. Falcón.

**Writing – original draft:** Sebastián Villada-Bedoya, Isaac G.-Santoyo.

**Writing – review & editing:** Sebastián Villada-Bedoya, Isaac G.-Santoyo, Aída Elizondo García, Osiris Gaona, Luisa I. Falcón.

**Competing interests:** The authors have declared that no competing interests exist.

## Supporting information

**S1 Table**. Biomarkers detected in Me’phaa children and adult populations. Differentially abundant clades among the age group and sex groups were identified by LEfSe using the highest LDA score. A total of 50 significantly different taxonomic biomarkers were detected.

(PDF File)

**S1 Fig**. Relative abundance of the GM and bacterial composition (16S rRNA V4) at the Family level in Me’phaa children and adults. Families with relative abundances < 1% were grouped into the “Others” category.

(PDF File)

